# Distribution of *luxS* and production of autoinducer-2 among gut Bacteroidales

**DOI:** 10.64898/2026.07.27.740950

**Authors:** Nolan W. Kennedy, Rebecca H. Gellman, Michael J. Coyne, Jessica C. Little, Ashley M. Sidebottom, Laurie E. Comstock

## Abstract

One of the best studied quorum sensing (QS) molecules, autoinducer-2 (AI-2), regulates processes in numerous bacteria. LuxS is an enzyme of the activated methyl cycle which, along with MtnN, converts *S*-adenosyl-homocysteine to homocysteine, releasing DPD (4,5-dihydroxy-2,3-pentanedione) which is spontaneously converted to AI-2. Many bacteria do not encode MtnN-LuxS and instead encode SahH, which directly converts *S*-adenosyl-homocysteine to homocysteine without AI-2 production. The genomes of some gut Bacteroidales were shown to contain *luxS*, however, these reports, as well as reports of the production of AI-2 by gut Bactereoidales have been inconsistent. We performed a comprehensive analysis of the distribution of *luxS* and *sahH* in Bacteroidota with an in-depth exploration of gut Bacteroidales. The data suggest that the ancestral Bacteroidota contained *sahH*, with numerous independent replacements with *mtnN-luxS* during diversification. In Bacteroidaceae, *Parabacteroides*, and many Prevotellaceae, *mtnN*-*luxS* or *sahH* are present in the same genetic region, adjacent to *yfhO*. Using *Bacteroides fragilis*, which contains *sahH,* and *Bacteroides uniformis* and *Phocaeicola vulgatus,* whose genomes contain *mtnN-luxS*, we show that *luxS*-containing strains produce AI-2. Transcriptomic analyses and gnotobiotic mouse experiments using wild-type strains and *mtnN-luxS* and *sahH* genetic swaps showed that Bacteroidaceae species do not respond to AI-2 under the conditions tested. However, analyses of 15 human gut metagenomic datasets show that *mtnN-luxS* containing Bacteroidales species average between 5.1 – 33% of the total bacteria in these various human gut microbiome datasets, and therefore, likely produce substantial amounts of AI-2 sensed by responsive gut symbionts and pathogens.

**Importance:** Here, we show that gut Bacteroidales possess one of two pathways to complete the activated methyl cycle, one that produces the QS molecule AI-2. We clarify conflicting data regarding AI-2 production in gut Bacteroidales and show that *luxS* is present in many gut Bacteroidales species, but lacking in species such as *Bacteroides thetaiotaomicron* and *Bacteroides fragilis*. We find that prevalent and abundant human gut Bacteroidales species including *Bacteroides uniformis* and *Phocaeicola vulgatus* produce substantial amounts of AI-2. While our data do not show that Bacteroidales sense or respond to AI-2 under the conditions tested, analyses of human gut metagenomic data reveal that AI-2 producing Bacteroidales comprise a large proportion of the gut bacteria of both industrialized and non-industrialized human populations.

## Introduction

A diverse gut microbiome is an important component of human health. Bacteroidales is one of the most stable and abundant orders of bacteria of the healthy human gut microbiota (1), and includes species of *Bacteroides*, *Phocaeicola*, *Parabacteroides* and *Segatella*, among others. Numerous Bacteroidales species simultaneously colonize the human gut at high density (2) where they frequently interact with each other. These interactions can be adversarial, as in the case of Type VI secretion system-mediated warfare (3), or beneficial, such as cross-feeding (4–6). Bacteroidales strains and species have different traits that enable these interactions.

How strains and species of Bacteroidales communicate with each other and other microbes through chemical signaling is less understood. The canonical process of interbacterial communication is quorum sensing (QS) where a secreted small molecule (Gram-negative) or secreted peptide (Gram-positive) induces a signaling cascade resulting in the expression of specific genes typically when the population reaches high cell density (7). This process coordinates and synchronizes group behaviors within the population such as biofilm formation, bioluminescence, and pathogenicity (8).

Two chemical signaling systems have recently been reported in Bacteroidota (9, 10). In these systems, small molecules are released from the bacteria where they can be sensed by kin or co-colonizing bacteria. These molecules accumulate and bind to transcriptional repressors to induce (de-repress) the expression of biosynthetic gene clusters (9) or type VI secretion systems (T6SS) and arrays of T6SS-specific immunity genes (10) on acquired immunity defense islands (11). Although these systems are analogous to QS systems, they regulate only a single or few genetic regions rather than having global gene regulatory effects.

While numerous small molecules act as QS signals, autoinducer-2 (AI-2) is described as a universal signaling molecule, in that it signals across species and kingdoms (12, 13). The gene responsible for AI-2 production, *luxS*, encodes an enzyme that is part of the activated methyl cycle (Fig. 1A)(14). The activated methyl cycle produces *S*-adenosylmethionine, which serves as the primary methyl donor for various methylation reactions. In species that have *luxS*, the gene *mtnN* (also known as *pfs*) is typically present, and their two products combine to convert *S*-adenosyl-homocysteine to homocysteine through an *S*-ribosylhomocysteine intermediate (Fig. 1A). During this conversion, DPD (4,5-dihydroxy-2,3-pentanedione) is released and spontaneously converted to AI-2 (15). When the AI-2 concentration is substantially high, it can either bind to a surface receptor and transmit a signal (16), or bind to an intracellular regulator to induce gene expression (17). In species that do not contain *mtnN-luxS*, the gene *sahH* is typically present. SahH directly converts *S*-adenosyl-cysteine to homocysteine to complete the activated methyl cycle, without production of DPD or AI-2 (Fig. 1A).

**Figure 1.**
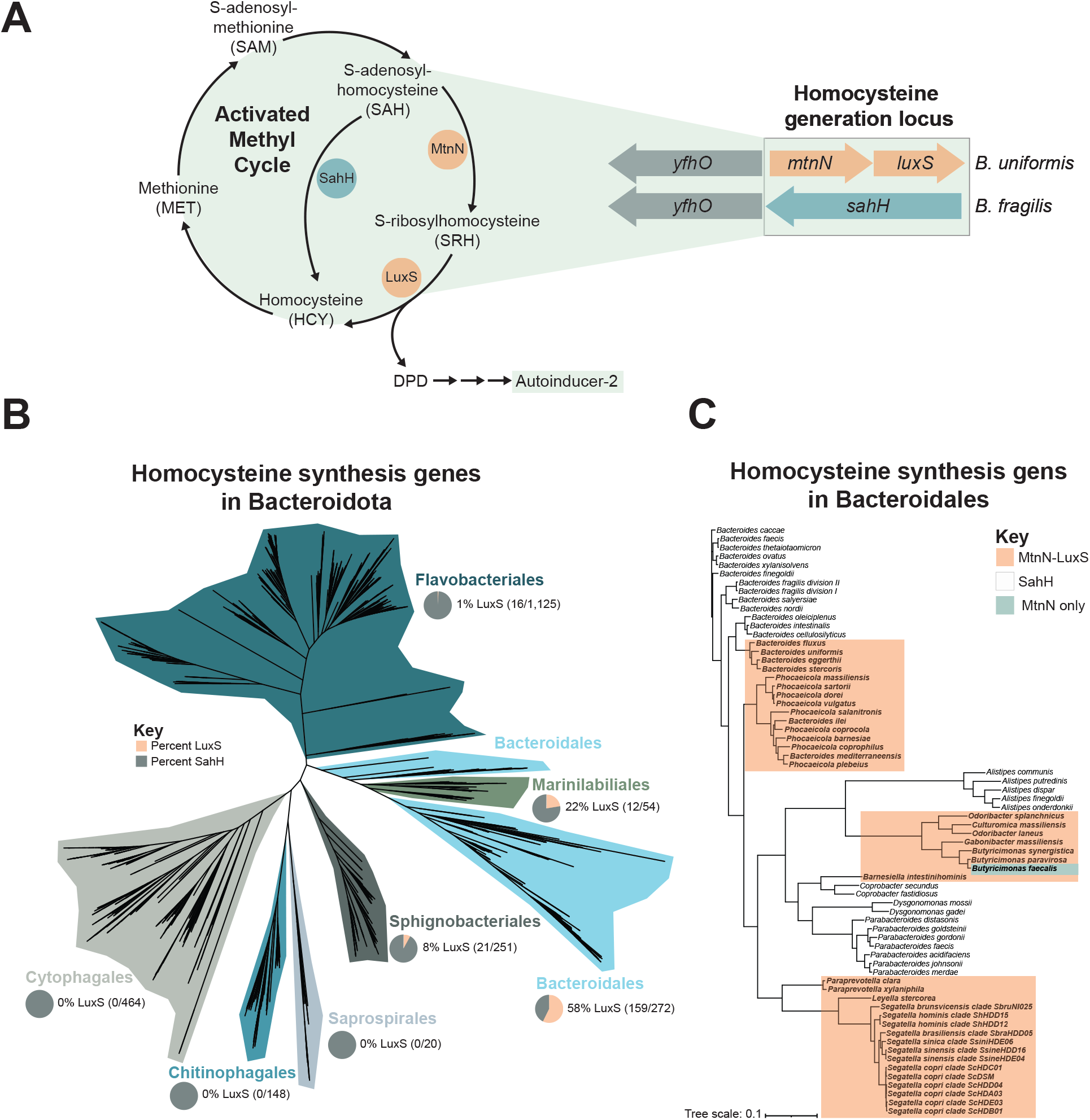
Characterization of homocysteine synthesis genes across Bacteroidota. A) Schematic representation of the activated methyl cycle (left) and homocysteine synthesis gene variability in Bacteroidota (right). Note that there are two possible routes for converting *S-*adenosylhomocysteine (SAH) to homocysteine (HCY), one single step route via the SahH protein that does not generate DPD and one two-step process involving the proteins MtnN and LuxS, which produces DPD as a byproduct. DPD is then spontaneously converted to AI-2 through multiple rearrangement steps. In Bacteroidota such as *B. uniformis*, the homocysteine generation locus contains genes for LuxS and MtnN, adjacent to the gene *yfhO*, while in Bacteroidota such as *B. fragilis*, only *sahH* is present. B) The abundance of MtnN-LuxS vs SahH at the homocysteine generation locus across Bacteroidota. Relative percentages in each clade are represented as a pie chart (light orange = MtnN-LuxS, grey = SahH). C) Phylogenetic tree of gut Bacteroidales showing those that encode MtnN-LuxS (orange background). Species with an uncolored background encode SahH.

Studies that investigated AI-2 production by gut Bacteroidales species date back more than 20 years and report conflicting findings. An early study (2005) showed that *B. fragilis*, *P. vulgatus* (then designated *B. vulgatus*) and *P. distasonis* (then designated *B. distasonis*), produce molecules that induce the *Vibrio campbellii* (previously designated *V. harveyi)* AI-2 reporter strain (18). Later, a study reported the PCR amplification of *luxS* genes from *B. fragilis* strain B3b and *P. vulgatus* ATCC 8482 and showed that these genes could restore AI-2 production to an *Escherichia coli luxS* mutant (19). A recent review presented a phylogenetic tree of 23 gut Bacteroidales strains showing that some possess *luxS* genes while others do not (20), but this study did not analyze many abundant human gut Bacteroidales species.

In the present study, we provide a comprehensive analysis of the presence of *mtnN-luxS* or *sahH* in sequenced Bacteroidota genomes, with a more in-depth focus of their presence in human gut Bacteroidales species. These analyses show that all strains of a species typically have either *mtnN*-l*uxS* or *sahH.* Despite previous work suggesting that *B. fragilis* possessed *luxS*, none of the 580 *B. fragilis* genomes analyzed contain *luxS*. To study the relevance of AI-2 production to gut Bacteroidales, we constructed genetic swaps in three prominent Bacteroidaceae species replacing *luxS-mtnN* with *sahH* or vice versa and performed transcriptomic analyses and mouse competitive colonization experiments.

## Results

### Distribution of *luxS* and *sahH* in Bacteroidota genomes

We searched 10,805 genomes of the phylum Bacteroidota, including all 7 orders, for orthologs of *luxS* and *sahH*. This investigation revealed that most genomes contain either *luxS* or *sahH*, but not both with only a few exceptions for endosymbiotic organisms and a few other outliers that lacked both (Table S1). In addition, of the genomes that contained either *luxS* or *sahH*, with few exceptions, strains of a given species contained either *luxS* or *sahH* revealing that these are core genes of a species.

This phylum-wide analysis showed that *sahH*-containing species dominate most orders. None of the genomes of the orders Cytophagales, Chitinophagales, and Saprospirales contain *luxS,* while nearly all contain *sahH* (464/469 species, 148/148 species, and 20/20 species, respectively) (Fig. 1B, Table S1). In the order Flavobacteriales, only 1.4% of species contain *luxS* (16 of 1141), most of the genus *Capnocytophaga* (Fig. 1B, Table S1). In Sphingobacteriales, of the 253 species queried, 21 have luxS.. Only two Bacteroidota orders have a high abundance of species that contain *luxS*: the Marinilabiliales in which 22.2% of species contain *luxS* (12 of 54) and the Bacteroidales where 55.4% of species contain *luxS* (159 of 287 species). These data strongly suggest that the ancestral Bacteroidota contained *sahH* with numerous independent replacements with *luxS* (or *mtnN-luxS*) at various branch points when families, genera and species diverged.

Closer inspection of human gut Bacteroidales revealed that there is a mix among species of the genus *Bacteroides* (Fig. 1C, Table S1). The genomes of many *Bacteroides* species such as *B. thetaoiotaomicron*, *B. ovatus*, and *B. fragilis* contain *sahH*, while *B. uniformis*, *B. eggerthii* and several other related species contain *luxS*. The genomes of *Parabacteroides*, and *Allistipes* species contain *sahH* whereas, *Phocaeicola, Paraprevotella,* and *Segatella* genomes contain *luxS*. At the species level, strains that have either *luxS* or *sahH* have the same gene, making them part of the core genome of a species.

### Genetic context of homocysteine biosynthesis genes

We analyzed the genes surrounding *mtnN-luxS* or *sahH* in the human gut Bacteroidales species to determine if these distinct genes are located in the same genomic site. We found that in *Bacteroides*, *Parabacteroides* and most *Phocaeicola* species, *mtnN-luxS* or *sahH* are in the same conserved genetic region where they are adjacent to a gene designated *yfhO*. (Fig 1A). *In silico* analyses leads us to predict that *yfhO* encodes the oligosaccharyltransferase (OTase) of the phylum-wide protein glycosylation system of Bacteroidota (21, 22). Despite the description of this general glycosylation system more than 15 years ago, the OTase that links the glycan chain to serine or threonine of the three amino acid glycosylation motif has yet to be identified (21, 23). There are four features of these YfhO proteins that suggest this protein family may encode the Bacteroidota OTase. The first is that HHPred analysis of the YfhO of *P. vulgatus* shows that an N-terminal portion of the protein aligns with the dolichyl-phosphate-mannose-protein mannosyltransferase of *Saccharomyces cerevisiae* (9E61, Probability: 98.08%, E-value: 0.00073) (24). Second, the protein glycosylation system is present in species of diverse orders of the phylum (22), and we show here that YfhO is largely conserved in the phylum (Table S1), absent in a minority of species including *Porphyromonas gingivalis*, some *Alistipes* species, and endosymbionts. Third, *yfhO* is predicted to be an essential gene of many Bacteroidales species (25–27) consistent with the finding that glycan truncations of the of the inner core are not tolerated (22). Lastly, YfhO, like all bacterial OTases, has numerous membrane-spanning domains, 12 predicated transmembrane regions for the *P. vulgatus* YfhO (Fig. S1A). Beyond *Bacteroides*, *Parabacteroides* and *Phocaeicola* species, there does not appear to be wide-spread genomic co-localization of *yfhO* with *mtnN-luxS* or *sahH*.

### Analysis of AI-2 production by gut Bacteroidales species

We sought to test the prediction that *mtnN-luxS* containing gut Bacteroidales strains produce AI- 2. We selected a subset of strains of abundant Bacteroidales species, four that possess *mtnN-luxS* (*P. vulgatus, Phocaeicola dorei, B. uniformis* and *Bacteroides eggerthii*) and five that possess *sahH* (*B. ovatus, B. thetaiotaomicron, B. fragilis, Parabacteroides distasonis,* and *Parabacteroides merdae*). We grew these representative species to near stationary phase (OD_600_ ∼1) in rich media and sampled the spent media to test for the presence of AI-2 using the *Vibrio campbelliii (*formerly *Vibrio harveyi*) reporter strain BB170 that luminesces in an AI-2 dose-dependent manner (28). AI-2 is typically at the highest extracellular concentration in late log or early stationary phase. Using this assay, we found that supernatants of species containing *mtnN-luxS* induced substantial luminescence (Figure 2A). None of the supernatants from *sahH*-containing species produced a signal greater than 1.5-times the negative control (blank media, Figure 2A). Notably, culture supernatants of *P. vulgatus* produced over 20 times greater signal than any of the *sahH*-containing species. These results indicated that *mtnN-luxS* containing Bacteroidales species produce and release AI-2.

**Figure 2.**
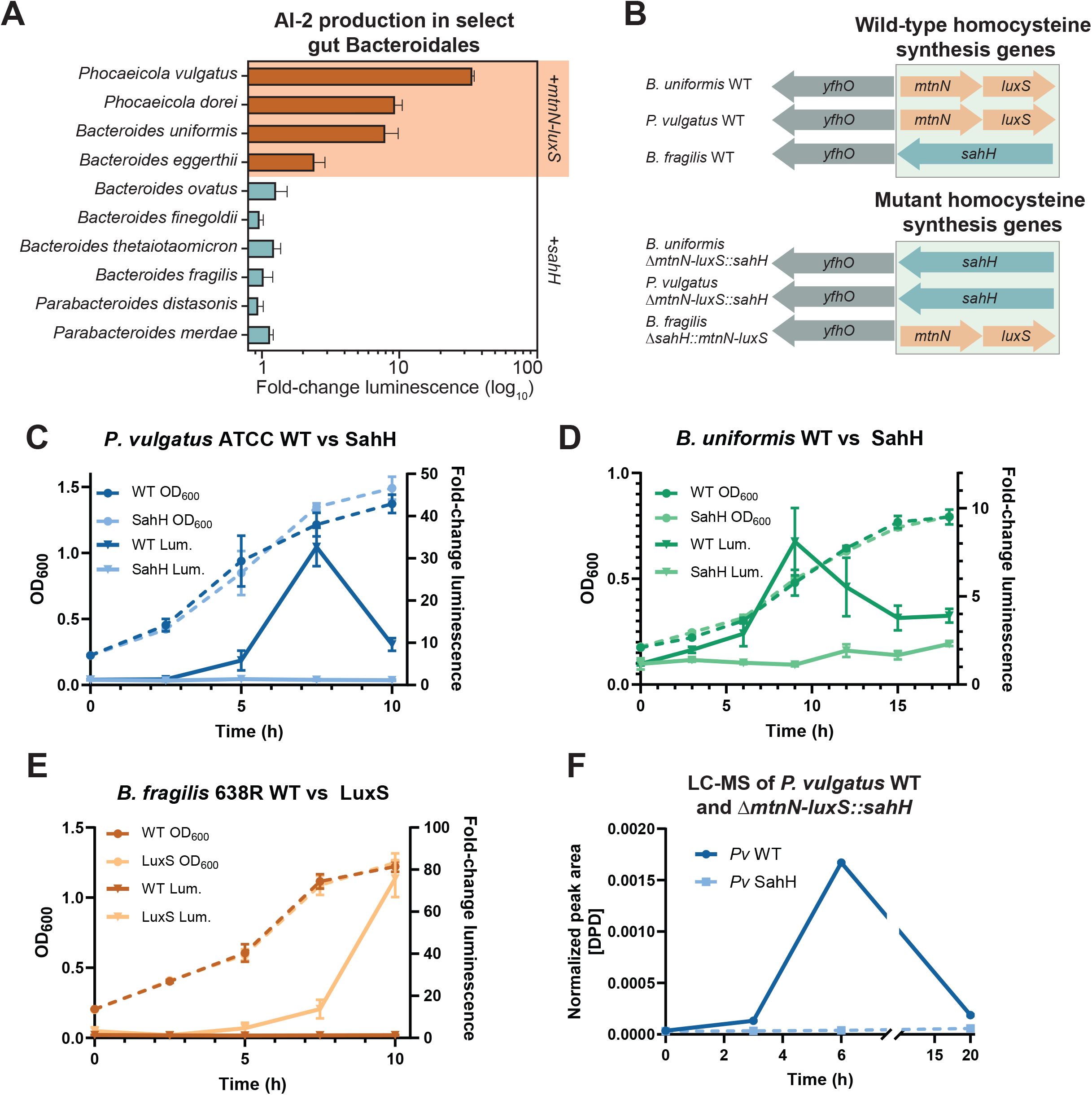
Gut Bacteroidales AI-2 production depends on *mtnN-luxS*. **A)** Select gut Bacteroidales species were tested for AI-2 production via the *V. campbellii* bioluminscence assay. Species containing *mtnN-luxS* are highlighted in orange with orange bars, species encoding *sahH* have blue bars. Results indicate log10 fold stimulation over the negative control condition (blank media). Error bars depict standard deviation of *n* = 3 biological replicates. **B)** Schematic representation of the homocysteine synthesis genes in three key gut Bacteroidales species. Wild-type loci are shown above, modified loci are shown below. **C-E)** Generation of AI-2 from WT and genetically modified gut Bacteroidales strains vs growth (OD_600_, left axes) over time. *P. vulgatus* WT and *sahH* (C, blue), *B. uniformis* WT and *sahH* (D, green), and *B. fragilis* WT and *mtnN-luxS* (E, orange) were grown, and spent media was sampled via the *V. campbellii* bioluminescence assay (normalized vs. negative control sample, right axis). Error bars show standard deviation of *n* = 3 biological replicates. **F)** LC-MS quantification (normalized relative to a blank media control) of DPD in spent media from *P. vulgatus* (*Pv* WT, dark blue) and *P. vulgatus* Δ*mtnN-luxS::sahH* (*Pv* SahH, light blue, dashed) over time. Error bars represent standard deviation of *n* = 3 biological replicates.

We next sought to abrogate AI-2 production to study the impact of the absence of AI-2 on these bacteria. Many studies aimed at eliminating AI-2 production in a bacterial strain simply delete *luxS* (29). However, this strategy produces a confounding variable – the disruption of the activated methyl cycle, which is an important metabolic process. To abrogate AI-2 production without disrupting the activated methyl cycle, we performed genetic swaps of *sahH* for *luxS-mtnN* and vice versa. We genetically modified strains of three species, two that contain *mtnN-luxS* (*P. vulgatus* and *B. uniformis*) and one that contains *sahH* (*B. fragilis*). In *B. uniformis* and *P. vulgatus*, the *mtnN-luxS* genes were swapped with *sahH*, generating a strain that should be deficient in AI-2 production while still maintaining a functional activated methyl cycle (Fig. 2B, Fig. S2). In *B. fragilis*, the native *sahH* gene was swapped for *mtnN-luxS* (Figure 2B), producing a strain that encodes the functions to produce AI-2 while maintaining a complete activated methyl cycle (Figure S2A).

We first tested the growth of the modified strains and found no growth defect in rich medium as all grew similarly to their wild-type (WT) counterparts (Fig.. S2B). Next, we tested if strains where *sahH* was swapped for *mtnN-luxS* produce AI-2. Cultures of WT and gene swapped strains were grown in rich medium and culture media were harvested at 2.5 or 3 h intervals (2.5 h for *P. vulgatus* and *B. fragilis,* 3 h for *B. uniformis*, which grew more slowly). In all three model species, strains encoding *mtnN-luxS*, whether naturally present or swapped in, had a noticeable spike in AI-2 production between mid-log or early stationary phase (Figure 2C-E). For WT *P. vulgatus*, AI-2 production began at 5 h and was most pronounced at 7.5 h, followed by a substantial drop in detectable AI-2 (Figure 2C). By comparison, the *P. vulgatus* Δ*mtnN-luxS*::*sahH* swapped strain produced no detectable AI-2. This drop may be due to the consumption and processing of AI-2 (30), and/or the acidification of the media at late growth phases, which was shown to inhibit the luminescence assay (31). To validate these results, liquid chromatography coupled with mass spectrometry (LC-MS) was used to confirm the presence of the AI-2 precursor, DPD, in the culture supernatants. As anticipated, DPD was highest in culture supernatants harvested from WT *P. vulgatus* at 6 h, followed by a drop in DPD (Fig. 2F). In the gene-swapped *P. vulgatus* Δ*mtnN-luxS*::*sahH*, no DPD was detected (Fig. 2F).

For *B. uniformis* and *B. fragilis*, similar trends were observed. In WT *B. uniformis*, AI-2 levels began to rise at 3 h, peaked at 9 h, then slowly dropped as the culture entered stationary phase (Fig 2D). Notably, the maximum level of AI-2 production in this strain was less than a third of the maximum level detected in the culture medium of WT *P. vulgatus* (Fig. 2C). By contrast, culture medium from the gene-swapped *B. uniformis* Δ*mtnN-luxS*::*sahH* strain induced minimal luminescence (Fig. 2D). For WT *B. fragilis*, which natively encodes *sahH*, no AI-2 was detected in the culture medium using the reporter strain (Fig 2E). However, the gene-swapped *B. fragilis* Δ*sahH::mtnN-luxS* culture medium showed the highest detected levels of AI-2 based on luminescence (Fig 2E). In this strain, AI-2 production was first detected in the culture medium at 5 h and peaked at 10 h. Together, these results show that SahH producing gut Bacteroidales strains can be engineered to produce AI-2 without detectable changes in growth.

### Analysis of response to AI-2

We next sought to determine if these species sense and respond to AI-2. We grew the non-AI-2-producing strains (WT *B. fragilis, P. vulgatus* Δ*mtnN-luxS*::*sahH,* and *B. uniformis* Δ*mtnN-luxS*::*sahH*) in the presence of 1 μM chemically synthesized DPD. This concentration was used as the linear range of the *V. campbellii* reporter assay is ∼0.4 µM to ∼35 µM, above which additional AI-2 becomes inhibitory (31). Additionally, we found that 1 µM AI-2 was at or above the amount of AI-2 found in spent cell cultures of our highest native AI-2 producing strains. After DPD addition, we assayed the culture media over time from these cultures to see if DPD, which spontaneously converts to AI-2, is taken up by cells. As DPD and AI-2 are labile, no AI-2 was detectable in the culture media after 20 h, even in media with added DPD without cells (Fig 3A). This finding validates the conclusion that the lack of detectable DPD in spent media via LC-MS after 20 hours is due to its instability (Fig 2F). After 6 h, there was less DPD in culture media from the *sahH*-encoding strains compared with uninoculated DPD containing media (Fig 3A).

**Figure 3.**
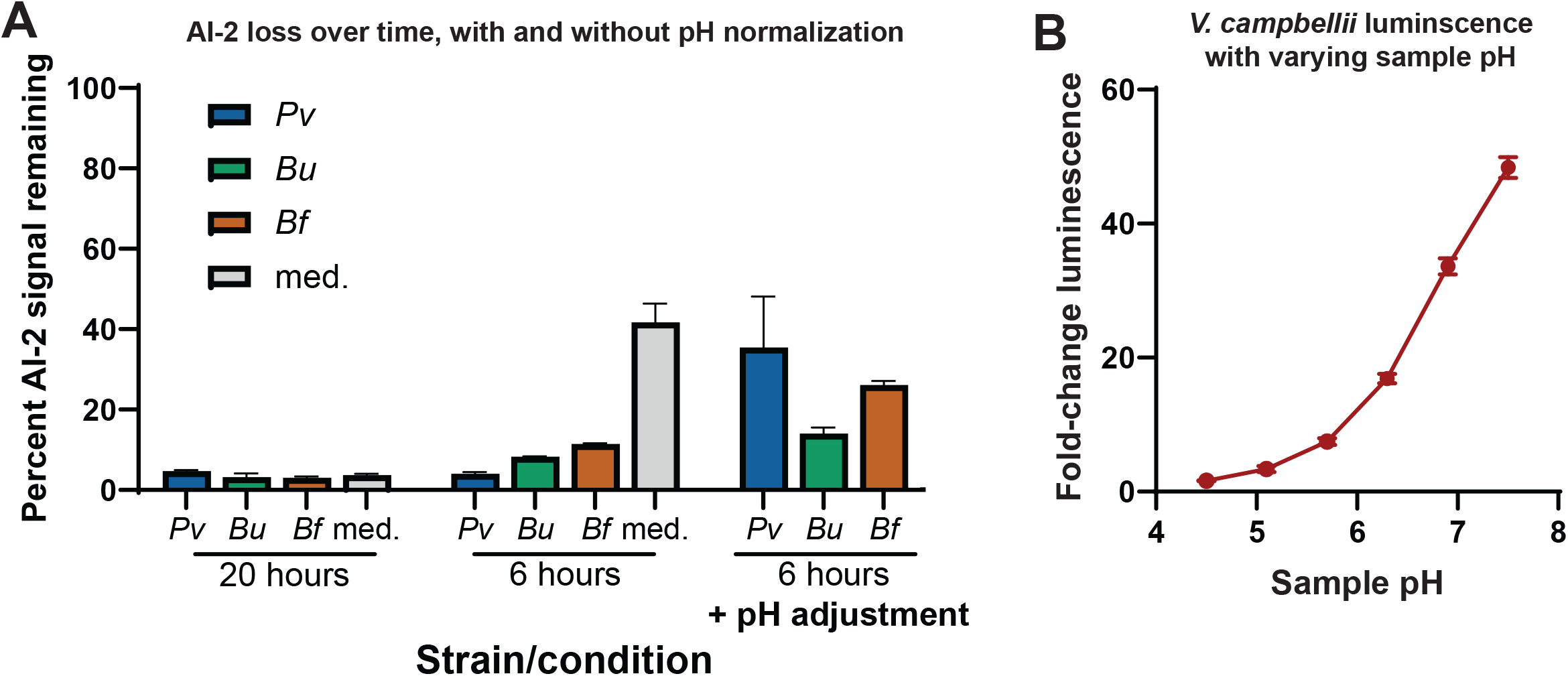
Loss of AI-2 over time in bacterial cultures. **A)** AI-2 consumption was measured for *P. vulgatus* (*Pv*, blue), *B. uniformis* (*Bu*, green), and *B. fragilis* (*Bf*, orange) cultures grown in the presence of 10 µM DPD, and the relative abundance vs a negative control (blank media) was measured after 6 hours and 20 hours post-subculture via the *V. campbellii* bioluminescence assay. Blank media without cells but with DPD was also tested to measure DPD loss over time. pH adjustment to neutral (pH = 7.0) for spent culture supernatant collected post-harvest (labelled “6 hours + pH adjustment”) partially recovered apparent DPD levels. Error bars represent standard deviation of *n* = 3 replicates. **B)** Low pH negatively impacts apparent levels of DPD as measured by the *V. campbellii* bioluminescence assay. Media with 1 µM DPD at various pH levels was tested using the *V. campbellii* bioluminescence assay (error bars represent standard deviation of *n* = 3 replicates).

However, as described in previous studies, acidification of the media at high cell density may interfere with the luminescence assay (31). Indeed, we found that acidified media decreased luminescence in the reporter assay when we tested a known concentration of DPD (10 µM) in media at various pH values (Fig 3B). To account for this, spent culture media from *sahH*-encoding strains were adjusted to a neutral pH (7.0) after harvest, which reduced the difference in luminescence between *sahH-*containing strains and DPD containing uninoculated media (Fig 3A). These results indicate that only minimal, if any, AI-2 is consumed by the Bacteroidales strains tested. However, lack of AI-2 consumption does not rule out that gut Bacteroidales can sense and respond to AI-2.

### Transcriptomic response to AI-2

To determine if Bacteroidales species can sense and respond to AI-2, we performed transcriptomic (RNASeq) analyses. Strains that natively encode SahH, or those that natively encode MtnN-LuxS but were engineered to produce SahH (WT *B. fragilis, P. vulgatus* Δ*mtnN-luxS*::*sahH,* and *B. uniformis* Δ*mtnN-luxS*::*sahH*) were grown in the presence of 1 μM chemically synthesized DPD. Two separate experiments were performed, in the first, DPD was added at the time of subculture, and in the second experiment DPD was added 1 h prior to harvest. In both experiments, cells were harvested for analysis at an OD600 of 0.7 when a quorum sensing response would be predicted to occur (Fig 2). These cultures were compared to the same strains grown with added sterile water (DPD solvent) instead of DPD. We detected no significant differences in gene expression in strains incubated with DPD compared to negative controls for any of the three strains in either experiment (no absolute value fold change ≥ 2.0 and no *p*-value < 0.05) (Tables S2, S3). Together, these results indicate that while MtnN-LuxS encoding gut Bacteroidales produce AI-2, they do not show a transcriptional response when DPD was added to the growth medium under the conditions of this assay.

Lastly, we transcriptionally profiled WT *P. vulgatus* and *P. vulgatus* with *sahH* swapped for *mtnN-luxS*. To determine if these strains have distinct gene expression profiles *in vivo*, we monoassociated gnotobiotic Swiss Webster mice with each strain. As the bacteria reach high density in the monocolonized mouse gut, we would expect to detect a response to AI-2 if such a capacity exists. After one week of colonization, fresh feces were collected and analyzed for differential gene expression between WT and the *sahH* variant using DESeq and EdgeR (Table S3). Other than *mtnN* and *luxS*, there were only two other genes that were significantly upregulated in the WT strain compared to the *sahH* strain by both statistical programs, YfhO (*p*adj 5.4 x 10^-5^) and BVU_2649 (*p*adj 9.4 x 10^-3^) encoding a protein of unknown function in a four gene operon likely involved in GDP-fucose synthesis. Two genes were downregulated in the WT *P. vulgatus* strain compared to the *sahH* replacements strain. One of these downregulated genes encodes a SusD outer surface protein (*p*adj 5.5 x 10^-4^), likely involved in nutrient import with its cognate SusC which just missed significance by the EdgeR program (*p*adj 2,4 x 10^-4^, FDR 0.054). The second gene just reached significance (*p*adj 0.029, FDR 0.05) and encodes a predicted diaminopimelate epimerase. The small number of differentially regulated genes, their level of significance and their putative encoded functions suggest these effects are unlikely due to differences in AI-2 production.

### *In vivo* fitness of WT and *mtnN-luxS/sahH* swapped Bacteroidales strains

We next investigated whether the *in vivo* fitness of gut Bacteroidales was altered when bacteria contained the heterologous gene(s) for homocysteine synthesis. We performed a series of competitive colonization assays in gnotobiotic mice (Figure 4A) wherein three male and three female germ-free Swiss Webster mice were inoculated with a 1:1 ratio of the *mtnN-luxS* and *sahH* isogenic variants of *B. fragilis, B. uniformis*, or *P. vulgatus*. The relative abundance of the two isogenic strains was measured at 14 days by qPCR of fecal samples, and fecal bacterial load was calculated by plating serial dilutions from each mouse (Figure 4B-D, S3). Total CFU/g reached an average of 2.3 x 10^10^ for *B. fragilis*, 2.8 x 10^10^ for *B. uniformis*, and 9.4 x 10^9^ for *P. vulgatus* (Fig. S3). Neither the *sahH* nor *mtnN-luxS* variants appeared to provide a fitness advantage to *B. fragilis*, *B. uniformis*, or *P. vulgatus* in both sets of mice in this short-term assay, though there was variability between the two groups of mice for each inoculum. The mean abundance of copies of *sahH* relative to copies of the housekeeping gene *gyrB* ranged from 8% to 76%. The difference in relative abundance did not correlate with the relative abundance of *mtnN-luxS* and *sahH* in the starting inoculum, or with the sex of the mice. Neither the WT nor mutant genotype (independent of whether the WT strain contained *sahH* or *luxS*) conferred a competitive advantage. Overall, the strains analyzed appeared equally fit in this short-term assay regardless of whether they retained their native gene(s) or had replacements with the heterologous gene(s).

**Figure 4.**
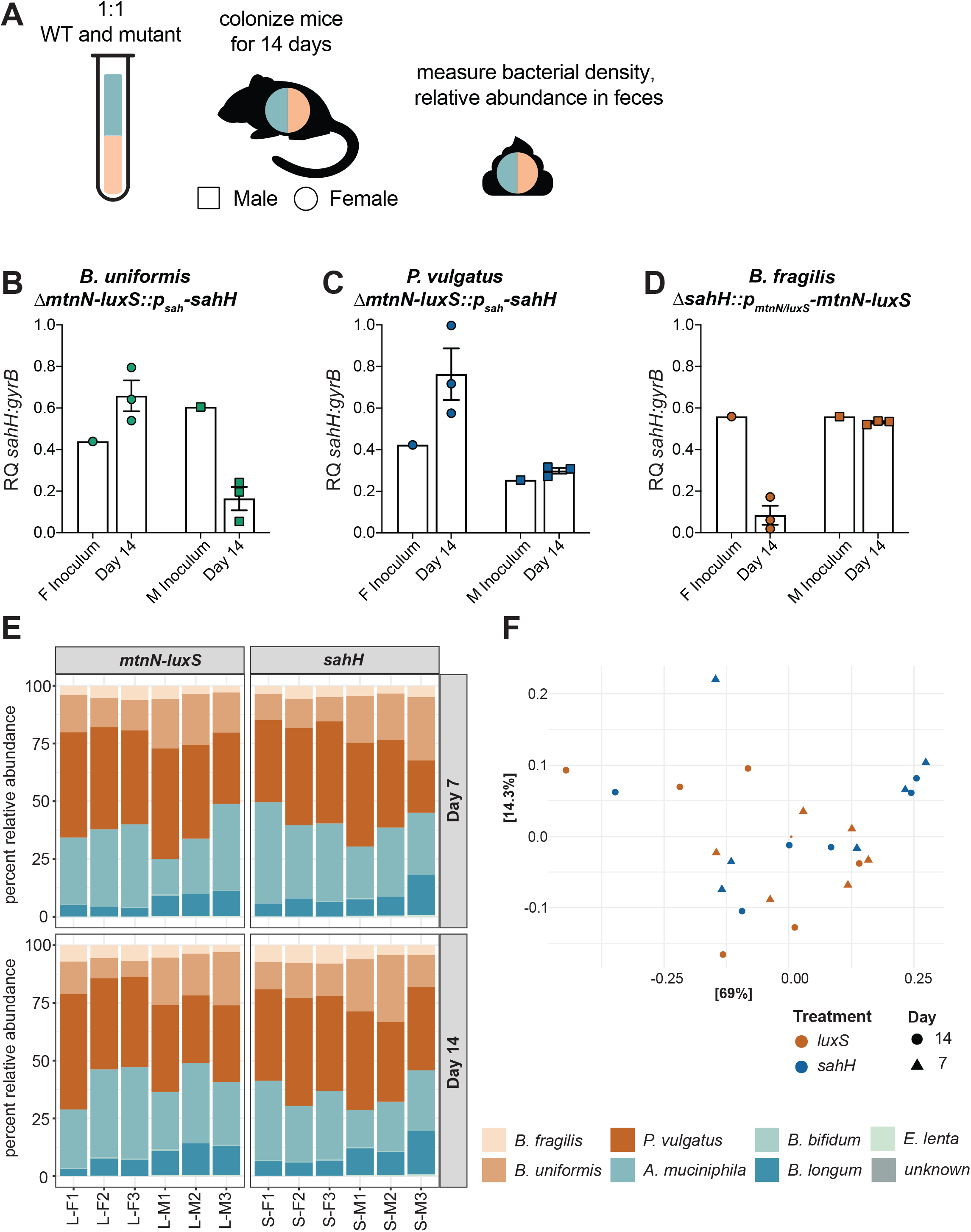
The presence of *mtnN-luxS* does not impact gut Bacteroidales fitness *in vivo.* **A)** Schematic of *in vivo* competition experiments (Swiss Webster mice, *n* = 3 mice per group, experiments repeated twice). **B-D)** Relative abundance of *sahH* compared to housekeeping gene (*gyrB*) measured by qPCR in inoculation culture and feces on Day 14 post-colonization (square indicates male mice, circle indicates female mice, error bars indicate SEM). Experiment performed with wild-type and *sahH/mtnN-luxS* swapped mutants of *B. uniformis* (B), *P. vulgatus* (C), and *B. fragilis* (D). **E)** Relative abundance of defined bacterial community containing Bacteroidales species with either *mtnN-luxS* or *sahH* in feces of gnotobiotic Swiss Webster mice on day 7 and 14 post-colonization, measured by 16S sequencing (*n* = 6 mice per group, F indicates female mouse, M indicates male mouse). **F)** PCoA plot of beta diversity (Bray-Curtis dissimilarity matrix) of in vivo bacterial communities shown in Fig. 4E. Color indicates treatment (blue = *sahH*, orange = *luxS*), shape indicates day post colonization. Each axis is defined by the dimensions that capture the greatest percentage of the variance of the data.

While AI-2 production did not appear to affect the relative fitness of single strains in competition assays, we tested whether the presence or absence of AI-2 would impact the composition of a more complex microbial community. We chose a set of four prevalent non-AI-2 producing human gut symbionts (*Akkermansia muciniphila, Bifidobacterium bifidum, Bifidobacterium longum,* and *Eggerthella lenta*) and colonized a set of female and male germ-free Swiss Webster mice with equal amounts of these strains, plus either all *sahH* or all *mtnN-luxS* isogenic variants of *B. fragilis*, *B. uniformis*, and *P. vulgatus*. The relative abundances of these seven-member communities were measured using 16S rRNA gene sequencing at days 7 and 14 post-gavage (Figure 4E). The Bacteroidales strains grew to a combined relative abundance of 63.3% and 58.5% in the AI-2 producing group and 59.39% and 64% in the non-producing group on days 7 and 14, respectively. *P. vulgatus* was the most abundant of the three Bacteroidales in both communities. Of the remaining community members, *A. muciniphila* and *B. longum* displayed relatively high colonization levels (average relative abundances of 28.5% and 9.5% on day 14, respectively), *B. bifidum* colonized minimally (<0.5% relative abundance), and *E. lenta* was not detectable in the samples, indicating it likely did not engraft. The beta diversity did not differ between the AI-2 producing and non-producing communities (Figure 4F), indicating that the production of AI-2 by these gut Bacteroidales species does not impact the structure of this small defined community over the course of this assay.

### Prevalence of Bacteroidales *luxS* and *sahH*-containing species in global gut microbiomes

Based on the human gut Bacteroidales species that encode MtnN-LuxS, we predicted that *luxS*-containing Bacteroidales would be highly prevalent and abundant in the gut microbiomes across human populations, where they likely contribute a substantial amount of AI-2 to the gut community. We analyzed 15 publicly available human gut metagenomic datasets including samples from 13 countries on 5 continents and Oceania (detailed in Table S4). Because each gut Bacteroidales species contains either *sahH* or *mtnN-luxS*, we measured the relative abundance and prevalence of each Bacteroidales species in these metagenomes to calculate the percent of total bacteria encoding a Bacteroidales LuxS or SahH. The combined relative abundance of *luxS*-containing Bacteoridales species in each population queried range from 5.13% (Sweden) to 33.03% (USA.Children), while the total relative abundance of *sahH*-containing Bacteroidales species in each population range from 0.41% (Fiji) to 12.63% (USA.Children), a much lower proportion of the microbial community in every dataset except Italy (Fig. 5A). In 8 of the 15 datasets, Bacteroidales species encoding LuxS are significantly more abundant than species encoding SahH (Fig. S4A, Welch’s t-test). This difference is driven by the highly abundant species *Phocaeicola vulgatus*, *Bacteroides uniformus*, and members of the genus *Segatella* (Fig. S4B). In only one dataset (Sweden) are *sahH*-containing Bacteroidales species significantly more prevalent than *luxS* containing Bacteroidales species (Fig. 5B). LuxS-containing species are significantly more prevalent in three datasets, whereas the remaining 11 datasets are not significantly different (Fig. S4C, D, Welch’s t-test). Together, these data indicate that LuxS-encoding Bacteroidales species comprise a considerable percentage of the total bacteria of human gut microbiomes across the globe. The AI-2 produced by these organisms likely impacts other members of the microbiota and enteric pathogens such as *Salmonella enterica*, which respond to AI-2 (30).

**Figure 5.**
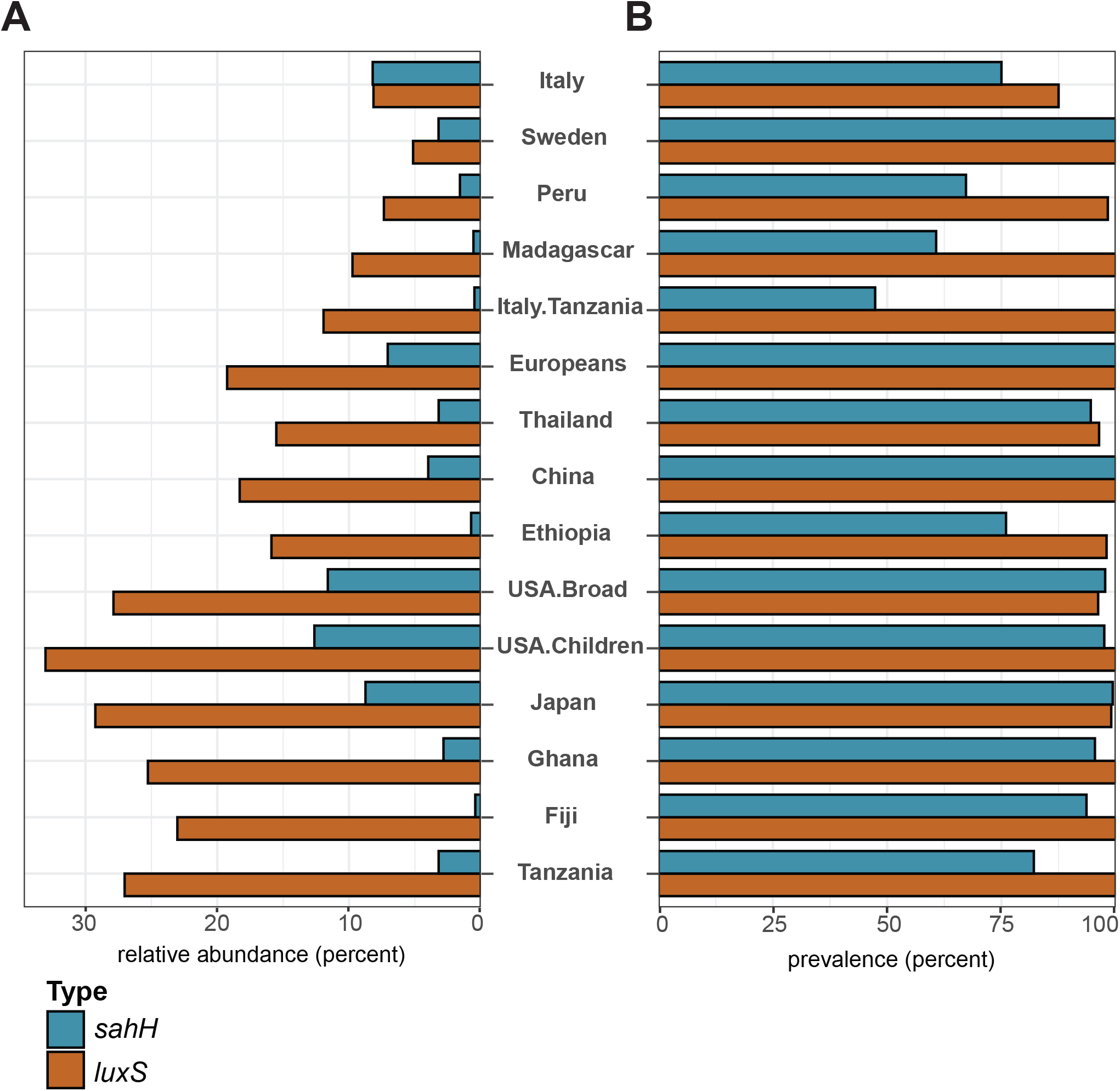
Variation of Bacteroidales *luxS* and *sahH* in global metagenomes. The total relative abundance **(A)** and prevalence **(B)** of *sahH-* or *luxS*-containing Bacteroidales species in metagenomic datasets of microbome data from studies of populations around the world. Color indicates whether the species possess *sahH* (blue) or *luxS* (orange).

## Discussion

Here we provide an in-depth analysis of *luxS* and AI-2 production in the human gut Bacteroidales, for which past studies have provided incomplete or inconsistent information. It was previously reported that *B. fragilis* and *P. distasonis* produce AI-2 (18, 19), which is inconsistent with the findings of our study. However, this work was performed prior to the ability to easily sequence genomes, and therefore these isolates were likely incorrectly speciated.

Based on the reported *luxS* gene sequences of these strains, we can now predict that they are likely *P. vulgatus*. Our examination of thousands of publicly available human gut Bacteroidales genomes indicates that *B. fragilis* and *P. distasonis* possess *sahH*, rather than *mtnN-luxS*. Additionally, our *in vitro* luminescence assays reveal that neither *B. fragilis* nor *P. distasonis* produce detectable amounts of AI-2.

A source of error in studies of AI-2 is that the typical bioluminescence assay that uses *V. campbellii* to detect the presence of AI-2 in bacterial supernatants is impacted by acidification of the culture medium (31). As shown in this study, when culture supernatants are not pH-normalized, the amount of AI-2 present in the sample can appear lower in this assay.

Another factor that can affect the apparent significance of AI-2 production to a bacterium is the use of *luxS* deletion mutants without accounting for effects on the disruption of the activated methyl cycle. In these studies, we replaced a native *Bacteroides mtnN-luxS* with a native *Bacteroides sahH* (and vice versa) to ensure that the activated methyl cycle was intact. A recent paper reported that the growth of *Clostridioides difficile* is inhibited by *B. fragilis*, but that this inhibition was lessened in a *luxS* mutant of *C. difficile* (29). Because this mutant cannot complete the activated methyl cycle, it is unclear if this effect is due to the absence of AI-2, or to some other change in *C. difficile* metabolism. Thus, future studies further investigating the role of AI-2 production in Bacteroidales or other microbes should utilize this swapped gene approach.

Despite the apparent lack of AI-2 sensing within Bacteroidales, it is likely that AI-2 produced by gut Bacteroidales plays a role in the gut microbiome community. It is possible that an AI-2 sensing non-Bacteroidales strain might respond to AI-2 made by Bacteroidales, given the high relative abundance of *luxS*-containing Bacteroidales in the gut. Furthermore, it is known that certain gut pathogens sense AI-2 leading to the induction of group behaviors. The presence or absence of Bacteroidales AI-2 could impact the quorum sensing, growth, or pathogenesis of one or more of these enteric pathogens. Previous work has shown that AI-2 is necessary for biofilm formation in the oral Bacteroidales species *Porphorymonas gingivalis*, regardless of whether *P. gingivalis* itself is producing AI-2 (32). The human commensal bacterium *Ruminococcus obeum* was reported to inhibit colonization of *V. cholerae* in the mouse gut, partially through AI-2 signaling (33). Using *E. coli* to increase the AI-2 levels in the mouse gut resulted in an increase in members of the Firmicutes (Bacillota) phylum, relative to the Bacteroidota (34). These studies, combined with the fact that other pathogens like *Salmonella enterica* sense and respond to AI-2, support that AI-2 produced by abundant gut Bacteroidales may impact gene expression in other gut bacteria.

What remains unknown is why species of human gut Bacteroidales use one of two genetically and enzymatically distinct paths to complete the activated methyl cycle, one of which produces the canonical quorum sensing molecule, and one that does not. Although our short-term competitive colonization experiment did not show that a strain is less fit if it produces the heterologous enzyme(s) for homocysteine synthesis, longer term experiments with a complex microbiota could reveal differences. The pattern of *luxS* presence within the Bacteroidales phylogenetic tree suggests that ancestral strains possessed *sahH,* and that *mtnN-luxS* was introduced later. Future work may uncover a benefit to maintaining this genetic variation within the human gut Bacteroidales.

## Methods

### Genome collections and phylogentic tree

An initial set of 14,243 Bacteroidota genomes downloaded from NCBI RefSeq on 02.27.25 was curated to remove any genome with incomplete taxonomy or other defects. The remaining 10,805 genomes (Table S1) comprises our Bacteroidota dataset. This dataset was parsed to select one genome to represent each species to build the phylogenetic tree. The strain chosen to represent the species was arrived at by first selecting RefSeq representative type strain entries, then eliminating remaining redundancy by selecting the genome with the largest N50. The resulting set of 2,373 genomes was used to create the tree (Fig, 1B), using VBCG (37) in RAxML mode(38). The output Newick file was used to create an unrooted tree using iToL(39). An additional collection of genomes was created to represent the human gut Bacteroidales (Table S1). This set was created by manual curation to retain Bacterodales isolates found in the Human gut, choosing representative isolates by examining the assembly level, assembly method, number of contigs, best N50, and other metrics. Duplicates of strains submitted by various repositories (ATCC, DSM, etc.) were removed, as were genome assemblies already identified as duplications in the literature (40, 41). Obvious longitudinal same-subject same-species collections were also parsed to retain the best representative using the criteria above. This curated collection comprising 3,429 human gut Bacteroidales genomes was further examined to correctly speciate *Segatella* isolates and differentiate them into clades by comparing the average nucleotide identity (ANI) of each of them to 13 Segatella genome sequences(42) using FastANI (43). *Bacteroides fragilis* isolates were identified as belonging to division I or II also using ANI measures in a similar manner.

### Phylogenetic distribution of LuxS, SahH, MtnN, and YfhO proteins

Each of the protein (faa) files from the 10,805 Bacteroidota genomes was searched (partial protein sequences were excluded) using hmmsearch for LuxS, SahH, MtnN, and YfhO using the gathering threshold cutoff for HMM profiles PF02664.19, PF05221.23, TIGR01704, and PF09586.17, respectively. The distribution of these proteins across various phylogenetic levels was then enumerated (Table S1).

### Bacterial culture

*Bacteroidales* strains were grown anaerobically at ∼37 °C. Broth cultures were inoculated from fresh brain heart infusion agar plates supplemented with 5 mg/liter hemin and 2.5 μg/liter vitamin (BHIS) into basal media (42). Cultures were grown to early-to-mid-log phase (OD_600_∼0.2-0.4) before subculturing in fresh supplemented basal media to an OD_600_ of 0.1 for experimentation. For cultures where the pH was adjusted to neutral, cells were grown anaerobically at 37 °C in broth as described above with a 10 mL culture volume in glass culture tubes. At harvest, cells were removed via centrifugation at 12,000 x *g* for 5 minutes. When indicated, sodium hydroxide was added to the supernatant dropwise to neutralize to a pH of ∼7.0, and AI-2 detection was carried out as described in the *V. campbellii* bioluminescence assay section. Antibiotics used for selection in this study included the following: carbenicillin (100 μg/mL), anhydrotetracycline (75 ng/mL), gentamicin (200 μg/mL), erythromycin (10 μg/mL), and cefoxitin (10 μg/mL). Details for the growth of *E. coli* for cloning and *V. campbellii* for AI-2 detection assays are described in their respective methods sections.

### Mutant construction

All primers used in this study are listed in Table S5. DNA fragments were PCR amplified with the primers listed in Table S5 with Phusion High-Fidelity DNA polymerase (New England Biolabs), then separated using gel electrophoresis and gel purified using a Monarch Spin DNA gel extraction kit (New England Biolabs). PCR products were ligated into the into the BamHI site of vectors and transformed into *E. coli* S17 λ pir and selected on Lysogeny Broth (LB) agar plates with 100 μg/mL (final concentration) carbenicillin. Clones were screened by PCR and sequence verified using whole-plasmid sequencing (Plasmidsaurus Inc., Louisville, KY).

For constructing genetically swapped mutants at the homocysteine generation locus, DNA segments flanking the gene(s) of interest (either *sahH* or *mtnN-luxS*) were PCR amplified using Phusion High-Fidelity DNA polymerase (New England Biolabs) using template DNA from their destination strain (either *B. fragilis* 638R, *P. vulgatus* ATCC 8482, or *B. uniformis* ATCC 8492).

Genes for insertion were amplified from their respective parent strains – *B. fragilis* 638R *(sahH*) or *P. vulgatus* ATCC 8482 (*mtnN-luxS*) – and inserted between flanking regions. DNA flanks and gene inserts were cloned into either pLGB13 (constructs to mate into *P. vulgatus* and *B. uniformis*) or pLGB36 (constructs to mate into *B. fragilis* 638R) (43, 44).

Plasmid constructs were transformed into *E. coli* S17 λ pir cells for conjugal transfer to target strains. Upon conjugation, Bacteroides cointegrates were selected on BHIS agar plates supplemented with 200 μg/mL gentamicin and 10 μg/mL cefoxitin. Cointegrate colonies were picked, cultured in supplemented basal media, grown for 3-5 h at ∼37 °C, and plated on BHIS agar plates supplemented with anhydrotetracycline. Double crossout recombinants were screened using colony PCR to confirm gene insertion.

### *V. campbellii* bioluminescence assay

Detection of AI-2 in culture supernatants was performed following the method developed by Taga and Xavier (45). Strains to be analyzed for AI-2 production were grown as described above in the “bacterial culture” section. For experiments where only endpoints were measured, cultures were grown to stationary phase. Samples from time course experiments were harvested at time points indicated in respective figures. For experiments performed in 96-well plates, 200 μL culture volumes were used; for experiments in test tubes, 5 mL culture volumes were used. Culture supernatants were harvested and centrifuged at 12,000 x *g* for 5 minutes. 190 μL of clarified supernatant was removed and stored at -80 °C until analysis.

The bioluminescence assay used in this study relies on the *V. campbellii* strain BB170, acquired from ATCC (BAA-1117), which luminesces in response to AI-2. For AI-2 detection experiments carried out in this study, BAA-1117 was inoculated from a 35% glycerol stock (stored at -80 °C) in 5 mL autoinducer bioassay (AB) media in a glass culture tube. AB media was prepared as previously described (45). This culture was grown at 30 °C with shaking at 250 rpm for ∼24 h.

In preparation for the bioluminescence assay, clarified supernatant samples to be analyzed were thawed slowly on ice. While samples thawed, positive and negative control samples were prepared. Chemically synthesized DPD (46) was purchased from ITQB NOVA (Instituto de Tecnologia Química e Biológica António Xavier, Portugal) and used at a final concentration of 1 μM diluted in AB media and served as the positive control, while blank AB media was used as a negative control. 10 μL of each control and clarified supernatant sample were pipetted into separate wells of a black, clear-bottom 96-well plate. Saturated BAA-1117 cultures were diluted 1:5000 into fresh AB media, and 90 μL of this diluted culture was pipetted into each well containing samples for analysis. The cultures were then incubated at 30 °C with continuous orbital shaking in a BioTek Epoch2 microplate spectrophotometer (Agilent Technologies, Santa Clara, CA),, measuring luminescence at 10 minute intervals. Data was analyzed as previously described (45). Experiments were performed in biological triplicate unless otherwise stated.

### LC-MS analysis of AI-2

*Bacteroides* cultures to be analyzed were grown as described in the “*In vitro* bacterial culture” section in glass culture tubes (4 mL culture volume). At indicated time points 600 μL of cell culture was harvested, centrifuged at 12,000 x *g* for 5 minutes, and flash frozen in a dry ice ethanol bath. Samples were stored at -80 °C until use.

Derivatization of DPD was adapted from the protocol described in Deng *et al.*(47). A 2,3-diaminonapthalene (DAN) was prepared by dissolving 10 mg DAN in 50 mL of 0.1 M HCl. An aliquot of water was used as a method blank and a pure standard of DPD was used for reference. Standard aliquots, quality control samples, or culture supernatants were mixed with an equal volume of DAN solution in 2 mL autosampler vials and vortexed on a thermomixer for 2 min to ensure thorough mixing.

Samples were then incubated at 90 °C for 40 min using a thermomixer. Following incubation, samples were allowed to cool to room temperature and subsequently centrifuged at 6000 rpm for 5 min at 20 °C. The supernatant was transferred to MS vial inserts, and 5 μL of each sample was injected onto the column for analysis.

Experimental and quality control samples were analyzed on a Thermo Fisher Vanquish liquid chromatography system coupled to an Orbitrap IQ-X mass spectrometer, operating in positive mode. 5 µL of sample was injected onto a Cortecs© UPLC T3 Column (1.2 µm, 2.1 x 100 mm) fitted with Cortecs© UPLC T3 guard at 50 °C. The mobile phase A was water with 5% acetonitrile and 0.1% formic acid and mobile phase B was acetonitrile with 0.1% formic acid. Gradient elution started with 0% B with a flow rate of 0.30 mL/min for 1.0 min and linearly increased to 100% B over 10 min and these conditions were held constant for 2.0 min. Finally, re-equilibration at 0% B was performed for 2 min. The electrospray ionization conditions were set with the spray voltage at 3.5 kV, vaporizer temp at 400 °C, ion transfer tube temp at 200 °C, and detection window set to 100-1000 mass-to-charge (*m/z)*.

Raw data files were analyzed in Thermo Freestyle software and peak lists were generated using MZmine 2.53(48). MZmine algorithms were applied to raw data for noise filtering, feature detection, and isotopic grouping. Noise was reduced by applying intensity thresholds, retaining only signals above 4E4 at the MS1 level and 1E4 at the MS2 level. Peak detection was performed using strict *m/z* and retention time tolerances (0.005 *m/z* and 0.10 min), and isotopologues grouped into single features. Peaks were normalized per sample to the total ion chromatogram by dividing the raw peak area of each compound by total signal in that sample. DPD-DAN product was identified using exact mass, retention time, and fragmentation pattern matching to the standard curve. Raw data files can be accessed under MassIVE accession number MSV000101223.

### Strain preparation for RNA-sequencing analysis

Strains grown for RNA sequencing were prepared as described in the “*In vitro* bacterial culture” section. After subculturing to an OD_600_ of 0.1 in fresh supplemented basal media, biological replicates of each strain were split into two separate culture tubes. For the experiment where DPD was added at subculture, one tube contained chemically synthesized DPD diluted in supplemented basal media to a final concentration of 1 μM. In the other tube, a volume of plain supplemented basal media equal to that of the diluted DPD was added as a negative control. For the experiment where DPD was added 1 h prior to harvest, biological replicate subcultures were still split into two separate tubes, but 1 μM DPD and plain supplemented basal media were added during the growth period rather than at subculture. In both experiments, cultures were grown anaerobically at 37 °C. OD_600_ was tracked using a Genesys 20 visible spectrophotometer (Thermo Fisher Scientific, Waltham, MA) and cells were harvested at an OD_600_ of ∼0.8. All strains were OD matched within 0.05 a.u. Cells were then harvested anaerobically, pelleted via centrifugation at 12,000 x *g* for 5 minutes, flash frozen in a dry ice ethanol bath, and stored at - 80 °C.

### RNASeq analysis

All samples for RNASeq were processed at the Duchossois Family Institute Microbiome Metagenomics Facility (MMF). Nucleic acid was recovered from bacterial pellets using the Maxwell RSC instrument (Promega Corporation, Madison, WI). RNA was quantified using a QubitFlex fluorometer (Life Technologies, Waltham, Massachusetts) and integrity was assessed using a Tapestation 4200 unit (Agilent Technologies, Inc., Santa Clara, CA). Ribosomal RNA was depleted from all samples using the NEBNext rRNA Bacterial and HMR depletion kits (New England Biolabs, Ipswich, MA). Libraries from ribosomal RNA-depleted samples were constructed using the Ultra Directional RNA library prep kit for Illumina (NEB), and normalized libraries of biological triplicates of all samples were sequenced on the NextSeq 2000 platform (Illumina, Inc., San Diego, CA) at 2 × 100 bp read length.BBMap (v. 38.90) was utilized for adapter and quality trimming of all reads, and processed reads were mapped to the appropriate bacterial genome sequence using the Bowtie 2 short-read aligner (v. 2.4.5) (49) and a General Feature Format file containing the intervals of protein-coding domains from the appropriate genome assembly. After conversion of the Bowtie 2 output to sorted and indexed BAM files using SAMtools (v. 1.11) (50), and BEDtools (v. 2.30.0) (51), the read mapping results were evaluated for differential gene expression using both DESeq2 (v. 1.48.1) (52) and edgeR (v. 4.6.2) (53), running under R (v. 4.5.2) (https://www.R-project.org). A gene was considered differentially expressed if both DESeq2 and edgeR detected that the absolute value of its expression level under experimental conditions differed from the control conditions by a fold change (FC) of ≥2 and by a p-value (padj for DESeq2 and FDR for edgeR) of ≤ 0.05. (Table S4).

### Gnotobiotic mouse experiments

All mouse experiments were approved by the Institutional Animal Care and Use Committee (IACUC), University of Chicago. All mice were bred at the Gnotobiotic Research Animal Facility (GRAF) at the University of Chicago. After inoculation, mice were housed in a cage rack system to maintain gnotobiotic status. All mice were germ-free Swiss Webster mice, and were 5-10 weeks old at the time of gavage.

For the competition assays, two competing strains were grown to equal ODs overnight and combined 1:1. Each mouse was gavaged with 200 µL of gavage culture. 1mL of the inoculation culture was pelleted and frozen at -80°C to measure the initial inoculum ratio. Feces were collected under sterile conditions on days 1 and 14 post-gavage. Fecal bacterial density was quantified by plating dilution series of samples from each mouse at each timepoint on BHI-S agar.

For the defined community experiments, individual overnight cultures of all bacterial strains were brought to the same OD_600_ and combined in equal amounts to make the inoculum. Each mouse was gavaged with 200 µL of the combined cultures. Feces were collected under sterile conditions on days 1, 7, and 14 post-gavage. DNA was extracted from both the gavage culture cell pellet and the mouse fecal samples using the QIAamp PowerFecal Pro DNA kit (Qiagen). Prior to extraction, samples were subjected to mechanical disruption using a bead beating method. Samples were briefly suspended in a bead tube (Qiagen) along with lysis buffer and loaded on a bead mill homogenizer (Qiagen, TissueLyser II). Samples were then centrifuged, and supernatant was resuspended in a reagent that effectively removes inhibitors. DNA was then purified using a silica spin column filter membrane and quantified using Qubit.

### qPCR

All DNA samples were normalized to 3 ng/uL. Quantification of *mtnN-luxS* or *sahH*-containing strains of *B. fragilis* 638R, *B. uniformis* ATCC 8492, and *P. vulgatus* ATCC 8482 was performed by designing qPCR primers against *B. fragilis* 638R *sahH*, and against each strain’s gyrase B gene (*gyrB*). Genomic DNA from each isolate was used to generate a standard curve for each primer pair. The efficiency value (E) for each primer pair was calculated as 10(1/−slope) of log10(DNA input) against Ct value. The proportion of *sahH* to *gyrB* was calculated using the ΔΔCt method.

### 16S rRNA gene sequencing

Samples were processed and sequenced by the DFI MMF. The V4-V5 region within the 16S ribosomal RNA gene was amplified using universal bacterial primers – 563F (5’-nnnnnnnn-NNNNNNNNNNNN-AYTGGGYDTAAA-GNG-3’) and 926R (5’-nnnnnnnn-NNNNNNNNNNNN- CCGTCAATTYHT-TTRAGT-3’), where ‘N’ represents the barcodes and ‘n’ are additional nucleotides added to offset primer sequencing. PCR conditions included initial denaturation at 94°C for 3 min followed by denaturation, annealing and extension at 94°C/15 s, 51°C/30 s and 72°C/1 min. Final extension was performed for 5 min. The approximately 360 bp amplicons were subsequently purified using a magnetic bead size selection. Purified amplicons were quantified using a QubitFlex Fluorometer (Invitrogen) and pooled at an equimolar concentration before Illumina compatible Combinatorial Dual Index (CDI) adapters were ligated using the QIAseq 1-step amplicon library kit (Qiagen). The completed libraries were sequenced on an Illumina MiSeq platform to generate 2x250bp paired-end reads.

### Analysis of 16S rRNA gene data

16S rRNA reads were processed using QIIME2 (v2024.5). Specifically, reads were demultiplexed with the q2-demux plugin after which denoising was performed using the DADA2 plugin. Reads were first trimmed at 190bp for both forward and reverse reads to remove low quality nucleotides. After trimming, reads with more than 2 erroneous bases were removed. Chimeras were detected and removed using DADA2’s consensus method. The remaining high quality ASVs had frequency recorded and were then classified on a Naive Bayes classifier trained on the RDP training data. Taxonomy of the ASVs were assigned to the species level using a custom database containing the genomes of all bacterial strains within the defined community with a minimum bootstrap confidence score of 80. Downstream analysis and graphing of 16S data was done in R (version 4.5.0).

### Analysis of global gut metagenomes for *luxS-*contianing Bacteroidales species

MetaPhlAn (v. 4.2.4, with database mpa_vJan25_CHOCOPhlAnSGB_202503) (54) was used to profile 15 metagenomic datasets (55–67). The resulting profiles were parsed for average relative abundance of species of the order Bacteroidales and prevalence of *luxS*- known containing species in these datasets (Table S4).

## Acknowledgments

We are grateful to Katia Flores for assistance with some genetic constructs. We thank R. Ramaswamy for initial project discussions. We thank Yessenia Sierra and the GRAF staff for gnotobiotic mouse breeding and maintenance. We thank V. Burgo and the DFI MMF staff for RNA sequencing. NWK was and RHG is supported by training grant T32DK007074 from NIH/NIDDK. This work was funded by the Duchossois Family Institute and grant R01AI093771 from the NIH/NIAID. The funders had no role in study design, data collection and interpretation, or the decision to submit the work for publication.

## References

1. Faith JJ, Guruge JL, Charbonneau M, Subramanian S, Seedorf H, Goodman AL, Clemente JC, Knight R, Heath AC, Leibel RL, Rosenbaum M, Gordon JI. 2013. The Long-Term Stability of the Human Gut Microbiota. Science 341:1237439.

2. Zitomersky NL, Coyne MJ, Comstock LE. 2011. Longitudinal Analysis of the Prevalence, Maintenance, and IgA Response to Species of the Order Bacteroidales in the Human Gut. Infect Immun 79:2012–2020.

3. Coyne MJ, Comstock LE. 2019. Type VI Secretion Systems and the Gut Microbiota. Microbiol Spectr 7.

4. Chen R, Zhan Z, Zhang S, Tang S, Qin H, Deng Z, Gao J. 2026. Duality of *Bacteroides* cross-feeding networks in health and disease. Pharmacol Res 230:108327.

5. Fernandez-Julia P, Commane DM, van Sinderen D, Munoz-Munoz J. 2022. Cross-feeding interactions between human gut commensals belonging to the Bacteroides and *Bifidobacterium* genera when grown on dietary glycans. Microbiome Res Rep 1:12.

6. Rakoff-Nahoum S, Coyne MJ, Comstock LE. 2014. An ecological network of polysaccharide utilization among human intestinal symbionts. Curr Biol CB 24:40–49.

7. Bassler BL. 1999. How bacteria talk to each other: regulation of gene expression by quorum sensing. Curr Opin Microbiol 2:582–587.

8. Papenfort K, Bassler B. 2016. Quorum-Sensing Signal-Response Systems in Gram-Negative Bacteria. Nat Rev Microbiol 14:576–588.

9. Linares-Otoya L, Shirkey JD, Chhetri BK, Mira A, Biswas A, Neff SL, Linares-Otoya MV, Chen Y, Campos-Florian JV, Ganoza-Yupanqui ML, Jeffrey PD, Hughson FM, Donia MS. 2025. Discovery of a widespread chemical signalling pathway in the Bacteroidota. Nature 646:423–432.

10. Tuzlak L, Pappas TE, Coyne MJ, Sheahan ML, Burgo V, Rice PA, Comstock LE. 2026. Antagonism by the type VI secretion system of *Bacteroides fragilis* is controlled by a TetR family regulator and released small molecule. Proc Natl Acad Sci U S A 123:e2516485123.

11. Ross BD, Verster AJ, Radey MC, Schmidtke DT, Pope CE, Hoffman LR, Hajjar AM, Peterson SB, Borenstein E, Mougous JD. 2019. Human gut bacteria contain acquired interbacterial defence systems. Nature 575:224–228.

12. Xavier KB, Bassler BL. 2003. LuxS quorum sensing: more than just a numbers game. Curr Opin Microbiol 6:191–197.

13. Miller MB, Bassler BL. 2001. Quorum sensing in bacteria. Annu Rev Microbiol 55:165–199.

14. Schauder S, Shokat K, Surette MG, Bassler BL. 2001. The LuxS family of bacterial autoinducers: biosynthesis of a novel quorum-sensing signal molecule. Mol Microbiol 41:463–476.

15. Chen X, Schauder S, Potier N, Van Dorsselaer A, Pelczer I, Bassler BL, Hughson FM. 2002. Structural identification of a bacterial quorum-sensing signal containing boron. Nature 415:545–549.

16. Zhang L, Li S, Liu X, Wang Z, Jiang M, Wang R, Xie L, Liu Q, Xie X, Shang D, Li M, Wei Z, Wang Y, Fan C, Luo Z-Q, Shen X. 2020. Sensing of autoinducer-2 by functionally distinct receptors in prokaryotes. Nat Commun 11:5371.

17. Li J, Attila C, Wang L, Wood TK, Valdes JJ, Bentley WE. 2007. Quorum Sensing in Escherichia coli Is Signaled by AI-2/LsrR: Effects on Small RNA and Biofilm Architecture. J Bacteriol 189:6011–6020.

18. Antunes LCM, Queiroz Ferreira L, Oliveira Ferreira E, Rodrigues Miranda K, Eliane Santos Avelar K, Maria Cavalcanti Pilotto Domingues R, Candida de Souza Ferreira M. 2005. *Bacteroides* species produce *Vibrio harveyi* autoinducer 2-related molecules. Anaerobe 11:295–301.

19. Peixoto RJM, Miranda KR, Ferreira EO, de Paula GR, Rocha ER, Lobo LA, Domingues RMCP. 2014. Production of AI-2 is mediated by the S-ribosylhomocystein lyase gene luxS in Bacteroides fragilis and *Bacteroides vulgatus*. J Basic Microbiol 54:644–649.

20. Oliveira RA, Cabral V, Torcato I, Xavier KB. 2023. Deciphering the quorum-sensing lexicon of the gut microbiota. Cell Host Microbe 31:500–512.

21. Fletcher CM, Coyne MJ, Villa OF, Chatzidaki-Livanis M, Comstock LE. 2009. A General O-Glycosylation System Important to the Physiology of a Major Human Intestinal Symbiont. Cell 137:321–331.

22. Coyne MJ, Fletcher CM, Chatzidaki-Livanis M, Posch G, Schaffer C, Comstock LE. 2013. Phylum-wide general protein O-glycosylation system of the Bacteroidetes. Mol Microbiol 88:772–783.

23. Plummer TH, Tarentino AL, Hauer CR. 1995. Novel, specific O-glycosylation of secreted *Flavobacterium meningosepticum* proteins. Asp-Ser and Asp-Thr-Thr consensus sites. J Biol Chem 270:13192–13196.

24. Du M, Yuan Z, Kovach A, Lyu M, Li H. 2025. Pmt4 recognizes two separate acceptor sites to O-mannosylate in the S/T-rich regions of substrate proteins. Nat Commun 16:9726.

25. Liu H, Shiver AL, Price MN, Carlson HK, Trotter VV, Chen Y, Escalante V, Ray J, Hern KE, Petzold CJ, Turnbaugh PJ, Huang KC, Arkin AP, Deutschbauer AM. 2021. Functional genetics of human gut commensal *Bacteroides thetaiotaomicron* reveals metabolic requirements for growth across environments. Cell Rep 34:108789.

26. Fiebig A, Schnizlein MK, Pena-Rivera S, Trigodet F, Dubey AA, Hennessy MK, Basu A, Pott S, Dalal S, Rubin D, Sogin ML, Eren AM, Chang EB, Crosson S. 2024. Bile acid fitness determinants of a Bacteroides fragilis isolate from a human pouchitis patient. mBio 15:e0283023.

27. Voogdt CGP, Roux I, Müller K, Karcher N, Bravo AM, Kalmar L, Varik V, Bobonis J, Zeller G, Zimmermann M, Patil KR, Typas A. 2025. A toolkit for transposon libraries and functional genomics in intestinal Bacteroidales. bioRxiv 10.1101/2025.10.10.681549.

28. Surette MG, Bassler BL. 1998. Quorum sensing in *Escherichia coli* and *Salmonella* typhimurium. Proc Natl Acad Sci 95:7046–7050.

29. Slater RT, Frost LR, Jossi SE, Millard AD, Unnikrishnan M. 2019. *Clostridioides difficile* LuxS mediates inter-bacterial interactions within biofilms. Sci Rep 9:9903.

30. Xavier KB, Bassler BL. 2005. Regulation of Uptake and Processing of the Quorum-Sensing Autoinducer AI-2 in *Escherichia coli*. J Bacteriol 187:238–248.

31. Vilchez R, Lemme A, Thiel V, Schulz S, Sztajer H, Wagner-Döbler I. 2007. Analysing traces of autoinducer-2 requires standardization of the *Vibrio harveyi* bioassay. Anal Bioanal Chem 387:489–496.

32. McNab R, Ford SK, El-Sabaeny A, Barbieri B, Cook GS, Lamont RJ. 2003. LuxS-Based Signaling in Streptococcus gordonii: Autoinducer 2 Controls Carbohydrate Metabolism and Biofilm Formation with *Porphyromonas gingivalis*. J Bacteriol 185:274–284.

33. Hsiao A, Shamsir Ahmed AM, Subramanian S, Griffin NW, Drewry LL, Petri WA, Haque R, Ahmed T, Gordon JI. 2014. Members of the human gut microbiota involved in recovery from *Vibrio cholerae* infection. Nature 515:423–426.

34. Thompson JA, Oliveira RA, Djukovic A, Ubeda C, Xavier KB. 2015. Manipulation of the Quorum Sensing Signal AI-2 Affects the Antibiotic-Treated Gut Microbiota. Cell Rep 10:1861–1871.

35. Tian R, Imanian B. 2023. VBCG: 20 validated bacterial core genes for phylogenomic analysis with high fidelity and resolution. Microbiome 11:247.

36. Stamatakis A. 2014. RAxML version 8: a tool for phylogenetic analysis and post-analysis of large phylogenies. Bioinformatics 30:1312–1313.

37. Letunic I, Bork P. 2024. Interactive Tree of Life (iTOL) v6: recent updates to the phylogenetic tree display and annotation tool. Nucleic Acids Res 52:W78–W82.

38. García-Bayona L, Coyne MJ, Comstock LE. 2021. Mobile Type VI secretion system loci of the gut Bacteroidales display extensive intra-ecosystem transfer, multi-species spread and geographical clustering. PLoS Genet 17:e1009541.

39. Zhang ZJ, Cole CG, Coyne MJ, Lin H, Dylla N, Smith RC, Pappas TE, Townson SA, Laliwala N, Waligurski E, Ramaswamy R, Woodson C, Burgo V, Little JC, Moran D, Rose A, McMillin M, McSpadden E, Sundararajan A, Sidebottom AM, Pamer EG, Comstock LE. 2024. Comprehensive analyses of a large human gut Bacteroidales culture collection reveal species- and strain-level diversity and evolution. Cell Host Microbe 32:1853–1867.e5.

40. Blanco-Míguez A, Gálvez EJC, Pasolli E, De Filippis F, Amend L, Huang KD, Manghi P, Lesker T-R, Riedel T, Cova L, Punčochář M, Thomas AM, Valles-Colomer M, Schober I, Hitch TCA, Clavel T, Berry SE, Davies R, Wolf J, Spector TD, Overmann J, Tett A, Ercolini D, Segata N, Strowig T. 2023. Extension of the *Segatella copri* complex to 13 species with distinct large extrachromosomal elements and associations with host conditions. Cell Host Microbe 31:1804–1819.e9.

41. Jain C, Rodriguez-R LM, Phillippy AM, Konstantinidis KT, Aluru S. 2018. High throughput ANI analysis of 90K prokaryotic genomes reveals clear species boundaries. 1. Nat Commun 9:5114.

42. Pantosti A, Tzianabos AO, Onderdonk AB, Kasper DL. 1991. Immunochemical characterization of two surface polysaccharides of *Bacteroides fragilis*. Infect Immun 59:2075–2082.

43. García-Bayona L, Comstock LE. 2019. Streamlined Genetic Manipulation of Diverse *Bacteroides* and *Parabacteroides* Isolates from the Human Gut Microbiota. mBio 10:e01762–19.

44. Ito T, Gallegos R, Matano LM, Butler NL, Hantman N, Kaili M, Coyne MJ, Comstock LE, Malamy MH, Barquera B. 2020. Genetic and Biochemical Analysis of Anaerobic Respiration in *Bacteroides fragilis* and Its Importance In Vivo. mBio 11:e03238–19.

45. Taga ME, Xavier KB. 2011. Methods for Analysis of Bacterial Autoinducer-2 Production. Curr Protoc Microbiol 23:1C.1.1–1C.1.15.

46. Ascenso OS, Marques JC, Santos AR, Xavier KB, Rita Ventura M, Maycock CD. 2011. An efficient synthesis of the precursor of AI-2, the signalling molecule for inter-species quorum sensing. Bioorg Med Chem 19:1236–1241.

47. Deng Z, Hou K, Valencak TG, Luo XM, Liu J, Wang H. 2022. AI-2/LuxS Quorum Sensing System Promotes Biofilm Formation of *Lactobacillus rhamnosus* GG and Enhances the Resistance to Enterotoxigenic Escherichia coli in Germ-Free Zebrafish. Microbiol Spectr 10:e0061022.

48. Pluskal T, Castillo S, Villar-Briones A, Orešič M. 2010. MZmine 2: Modular framework for processing, visualizing, and analyzing mass spectrometry-based molecular profile data. BMC Bioinformatics 11:395.

49. Langmead B, Salzberg SL. 2012. Fast gapped-read alignment with Bowtie 2. 4. Nat Methods 9:357–359.

50. Danecek P, Bonfield JK, Liddle J, Marshall J, Ohan V, Pollard MO, Whitwham A, Keane T, McCarthy SA, Davies RM, Li H. 2021. Twelve years of SAMtools and BCFtools. GigaScience 10:giab008.

51. Quinlan AR, Hall IM. 2010. BEDTools: a flexible suite of utilities for comparing genomic features. Bioinformatics 26:841–842.

52. Love MI, Huber W, Anders S. 2014. Moderated estimation of fold change and dispersion for RNA-seq data with DESeq2. Genome Biol 15:550.

53. Robinson MD, McCarthy DJ, Smyth GK. 2010. edgeR: a Bioconductor package for differential expression analysis of digital gene expression data. Bioinformatics 26:139–140.

54. Blanco-Míguez A, Beghini F, Cumbo F, McIver LJ, Thompson KN, Zolfo M, Manghi P, Dubois L, Huang KD, Thomas AM, Nickols WA, Piccinno G, Piperni E, Punčochář M, Valles-Colomer M, Tett A, Giordano F, Davies R, Wolf J, Berry SE, Spector TD, Franzosa EA, Pasolli E, Asnicar F, Huttenhower C, Segata N. 2023. Extending and improving metagenomic taxonomic profiling with uncharacterized species using MetaPhlAn 4. Nat Biotechnol 41:1633–1644.

55. Yachida S, Mizutani S, Shiroma H, Shiba S, Nakajima T, Sakamoto T, Watanabe H, Masuda K, Nishimoto Y, Kubo M, Hosoda F, Rokutan H, Matsumoto M, Takamaru H, Yamada M, Matsuda T, Iwasaki M, Yamaji T, Yachida T, Soga T, Kurokawa K, Toyoda A, Ogura Y, Hayashi T, Hatakeyama M, Nakagama H, Saito Y, Fukuda S, Shibata T, Yamada T. 2019. Metagenomic and metabolomic analyses reveal distinct stage-specific phenotypes of the gut microbiota in colorectal cancer. Nat Med 25:968–976.

56. Lloyd-Price J, Arze C, Ananthakrishnan AN, Schirmer M, Avila-Pacheco J, Poon TW, Andrews E, Ajami NJ, Bonham KS, Brislawn CJ, Casero D, Courtney H, Gonzalez A, Graeber TG, Hall AB, Lake K, Landers CJ, Mallick H, Plichta DR, Prasad M, Rahnavard G, Sauk J, Shungin D, Vázquez-Baeza Y, White RA, Braun J, Denson LA, Jansson JK, Knight R, Kugathasan S, McGovern DPB, Petrosino JF, Stappenbeck TS, Winter HS, Clish CB, Franzosa EA, Vlamakis H, Xavier RJ, Huttenhower C. 2019. Multi-omics of the gut microbial ecosystem in inflammatory bowel diseases. Nature 569:655–662.

57. Karlsson FH, Tremaroli V, Nookaew I, Bergström G, Behre CJ, Fagerberg B, Nielsen J, Bäckhed F. 2013. Gut metagenome in European women with normal, impaired and diabetic glucose control. Nature 498:99–103.

58. Qin J, Li R, Raes J, Arumugam M, Burgdorf KS, Manichanh C, Nielsen T, Pons N, Levenez F, Yamada T, Mende DR, Li J, Xu J, Li S, Li D, Cao J, Wang B, Liang H, Zheng H, Xie Y, Tap J, Lepage P, Bertalan M, Batto J-M, Hansen T, Le Paslier D, Linneberg A, Nielsen HB, Pelletier E, Renault P, Sicheritz-Ponten T, Turner K, Zhu H, Yu C, Li S, Jian M, Zhou Y, Li Y, Zhang X, Li S, Qin N, Yang H, Wang J, Brunak S, Doré J, Guarner F, Kristiansen K, Pedersen O, Parkhill J, Weissenbach J, Bork P, Ehrlich SD, Wang J, Blottiere H, Borruel N, Bruls T, Casellas F, Chervaux C, Cultrone A, Delorme C, Denariaz G, Dervyn R, Forte M, Friss C, Guchte M van de, Guedon E, Haimet F, Jamet A, Juste C, Kaci G, Kleerebezem M, Knol J, Kristensen M, Layec S, Roux KL, Leclerc M, Maguin E, Minardi RM, Oozeer R, Rescigno M, Sanchez N, Tims S, Torrejon T, Varela E, Vos W de, Winogradsky Y, Zoetendal E, Bork P, Ehrlich SD, Wang J. 2010. A human gut microbial gene catalogue established by metagenomic sequencing. Nature 464:59–65.

59. Liu W, Zhang J, Wu C, Cai S, Huang W, Chen J, XI X, Liang Z, Hou Q, Zhou B, Qin N, Zhang H. 2016. Unique Features of Ethnic Mongolian Gut Microbiome revealed by metagenomic analysis. Sci Rep 6:34826.

60. Flannery JE, Stagaman K, Burns AR, Hickey RJ, Roos LE, Giuliano RJ, Fisher PA, Sharpton TJ. 2020. Gut Feelings Begin in Childhood: the Gut Metagenome Correlates with Early Environment, Caregiving, and Behavior. mBio 11:e02780–19.

61. Brito I, Yilmaz S, Huang K, Xu L, Jupiter S, Jenkins A, Naisilisili W, Tamminen M, Smillie C, Wortman J, Birren B, Xavier R, Blainey P, Singh A, Gevers D, Alm E. 2016. Mobile genes in the human microbiome are structured from global to individual scales. Nature 535:435–439.

62. Tett A, Huang KD, Asnicar F, Fehlner-Peach H, Pasolli E, Karcher N, Armanini F, Manghi P, Bonham K, Zolfo M, De Filippis F, Magnabosco C, Bonneau R, Lusingu J, Amuasi J, Reinhard K, Rattei T, Boulund F, Engstrand L, Zink A, Collado MC, Littman DR, Eibach D, Ercolini D, Rota-Stabelli O, Huttenhower C, Maixner F, Segata N. 2019. The *Prevotella copri* Complex Comprises Four Distinct Clades Underrepresented in Westernized Populations. Cell Host Microbe 26:666–679.e7.

63. Vangay P, Johnson AJ, Ward TL, Al-Ghalith GA, Shields-Cutler RR, Hillmann BM, Lucas SK, Beura LK, Thompson EA, Till LM, Batres R, Paw B, Pergament SL, Saenyakul P, Xiong M, Kim AD, Kim G, Masopust D, Martens EC, Angkurawaranon C, McGready R, Kashyap PC, Culhane-Pera KA, Knights D. 2018. U.S. immigration westernizes the human gut microbiome. Cell 175:962–972.e10.

64. Obregon-Tito AJ, Tito RY, Metcalf J, Sankaranarayanan K, Clemente JC, Ursell LK, Zech Xu Z, Van Treuren W, Knight R, Gaffney PM, Spicer P, Lawson P, Marin-Reyes L, Trujillo-Villarroel O, Foster M, Guija-Poma E, Troncoso-Corzo L, Warinner C, Ozga AT, Lewis CM. 2015. Subsistence strategies in traditional societies distinguish gut microbiomes. Nat Commun 6:6505.

65. Pasolli E, Asnicar F, Manara S, Zolfo M, Karcher N, Armanini F, Beghini F, Manghi P, Tett A, Ghensi P, Collado MC, Rice BL, DuLong C, Morgan XC, Golden CD, Quince C, Huttenhower C, Segata N. 2019. Extensive Unexplored Human Microbiome Diversity Revealed by Over 150,000 Genomes from Metagenomes Spanning Age, Geography, and Lifestyle. Cell 176:649–662.e20.

66. Rampelli S, Schnorr SL, Consolandi C, Turroni S, Severgnini M, Peano C, Brigidi P, Crittenden AN, Henry AG, Candela M. 2015. Metagenome Sequencing of the Hadza Hunter-Gatherer Gut Microbiota. Curr Biol 25:1682–1693.

67. D’Amico F, Soverini M, Zama D, Consolandi C, Severgnini M, Prete A, Pession A, Barone M, Turroni S, Biagi E, Brigidi P, Masetti R, Rampelli S, Candela M. 2019. Gut resistome plasticity in pediatric patients undergoing hematopoietic stem cell transplantation. Sci Rep 9:5649.

